# A Computational Functional Tissue Unit of the Human Myometrium for *In Silico* Study of Gestational Excitability and Pathophysiology

**DOI:** 10.64898/2026.05.05.723096

**Authors:** Jagir R. Hussan, Shawn A. Means, Peter J. Hunter, Alys R. Clark

**Affiliations:** Auckland Bioengineering Institute, University of Auckland, Auckland, New Zealand

**Keywords:** myometrium, computational modelling, electrophysiology, emergent pacemakers

## Abstract

The human myometrium undergoes a dramatic transformation during pregnancy, shifting from quiescence to highly synchronised contractility. Understanding this transition is crucial for addressing pathologies such as preterm labour and dystocia (ineffective labour). We present a multi-scale Functional Tissue Unit (FTU) model allowing us to investigate how tissue-level excitability emerges from single-cell electrophysiology. We propose a heterogeneity-driven selection mechanism, wherein a sub-population of cells with high intrinsic excitability dynamically emerges as pace-makers. This active process complements passive depolarisation by interstitial cells, allowing spontaneous excitation to arise without a fixed anatomical pacemaker. Stochastic simulations produced an average burst frequency of 0.047 Hz (*≈*2.8 bursts per minute), closely consistent with clinical measurements of 2–3 contractions per minute during active labour, and demonstrated that this function is robust to spatial topological changes. Furthermore, implementation of inflammation-induced remodelling simulations successfully linked molecular-level changes to a preterm labour phenotype. This model provides a platform for investigating uterine contractility and serves as a component for future whole-organ Physiome models.

## Introduction

The transition of the myometrium, the uterine smooth muscle, from relative quiescence to high excitability is critical for successful delivery; dysregulation of this process leads to complications such as preterm labour and dystocia Wray and Arrowsmith (2021); Smith (2007). Unlike the heart, the uterus lacks a defined anatomical pacemaker Lammers (2013); Garrett et al. (2022). Rhythmicity is currently thought to arise from either passive depolarisation by interstitial cells (e.g., telocytes Duquette et al. (2005); Means et al. (2025); Hutchings et al. (2009)) or intrinsic mechanisms within uterine smooth muscle cells (USMCs). Here, we focus on the latter, testing the hypothesis that pacemaker identity can emerge dynamically from cellular heterogeneity and the gestational downregulation of K^+^ channels (the ‘potassium switch’ Greenwood and Tribe (2014); Brainard et al. (2007); Khan et al. (1993)).

To investigate these temporal and spatial dynamics, mathematical modelling has increasingly provided critical insights into myometrial contractility across multiple scales Garrett et al. (2022). At the cellular level, comprehensive biophysical frameworks were developed to capture the complex dynamics of transmembrane currents and calcium handling Rihana et al. (2009); Tong et al. (2011). Expanding to the syncytial level, the foundational work by Sheldon *et al*. highlighted the pivotal role of spatial heterogeneity, demonstrating how stochastic variations in cellular coupling Sheldon et al. (2013) and dynamic alterations in gap junction connexin ratios Sheldon et al. (2014) dictate the transition from local quiescence to global excitation. Building upon this, the most recent modelling efforts have advanced towards fully three-dimensional, multiscale, and anisotropic representations of the uterus, integrating specific fibre architectures to track electrical propagation from the cellular source to the abdominal surface Yang et al. (2024); Means et al. (2025). While these macro-scale models excel at exploring global wave propagation and surface electrohysterography, there remains a need to isolate the local micro-architecture.

Therefore, by isolating this active cellular mechanism within a multi-scale Functional Tissue Unit (FTU) framework de Bono (2012); Hunter and Nielsen (2005), we assess whether intrinsic USMC remodelling alone is sufficient to generate coordinated contractions, essentially providing the myometrium with a robust mechanism for rhythmicity independent of interstitial cells Xu et al. (2015). Our model identifies USMCs with naturally high T-type Ca^2+^ conductance (*g*_*CaT*_) Lee et al. (2009) and applies a targeted reduction in repolarising K^+^ currents (*g*_*Kvm*_). This targeted remodelling allows these cells to overcome the hyperpolarising current sink—or quiescence-clamp imposed by the surrounding passive cell network that otherwise restrains intrinsic excitation Sheldon et al. (2013). This approach enables us to investigate how spontaneous, propagating waves of excitation emerge from a heterogeneous cell population driven by gestationally regulated ion channel remodelling.

## Methods

### A. The Single Smooth Muscle Cell Model

The core of the Functional Tissue Unit (FTU) is a biophysically detailed model of the Uterine Smooth Muscle Cell (USMC). We selected the comprehensive biophysical formulation by Tong *et al*. Tong et al. (2011) over more recent reduced models (e.g., Means et al. (2023)). Because our methodology relies heavily on non-dimensional stiffness analysis to explicitly isolate distinct physiological timescales, preserving the full transient, non-steady-state dynamics of the gating variables was strictly necessary.

To account for intracellular calcium handling, which is critical for the refractory period and burst termination, we extended the original model to include Sarcoplasmic Reticulum (SR) dynamics incorporating SERCA pumps and ryanodine receptors, as described in previous work Means et al. (2025); Matsuki et al. (2017); Garrett et al. (2022).

The membrane potential *V*_*m*_ is governed by the conservation of charge:

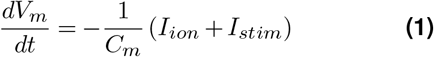

where *C*_*m*_ is the membrane capacitance (pF), *I*_*ion*_ is the sum of transmembrane ionic currents (pA), and *I*_*stim*_ is the external stimulus current (pA).

The gating variables (*x*) for the ion channels follow standard Hodgkin–Huxley kinetics:

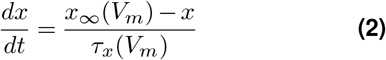

where *x*_*∞*_ represents the steady-state activation/inactivation and *τ*_*x*_ is the voltage-dependent relaxation time constant. See Figure 1 for a schematic of the cell model components included in *I*_*ion*_ and the SR Ca^2+^dynamics.

**Figure 1.**
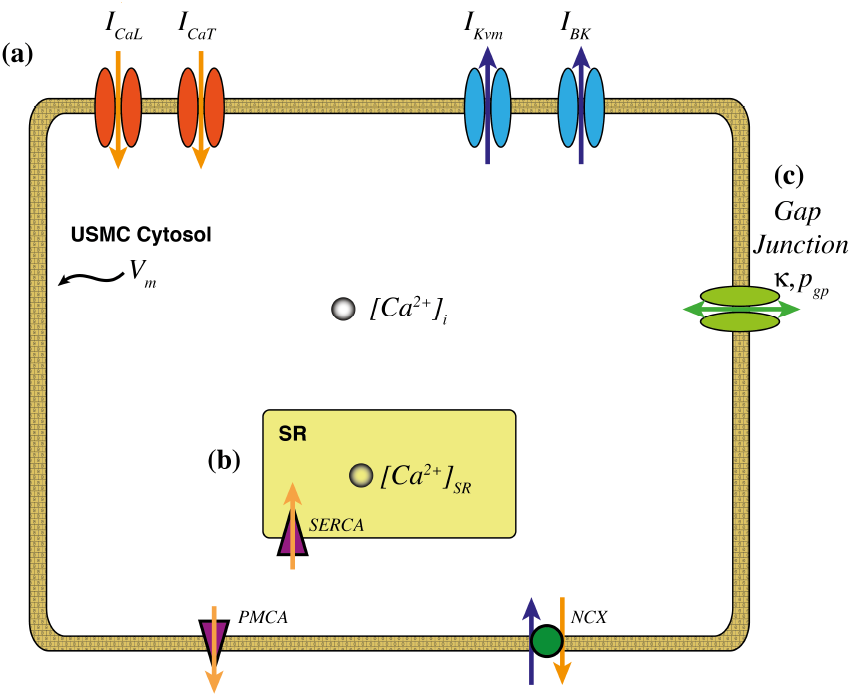
Schematic of the single-cell USMC model components. **(a)** Key transmembrane currents include the L-type (*I*_*CaL*_) and T-type (*I*_*CaT*_) calcium channels, and the voltage-gated potassium channels (*I*_*Kvm*_) involved in the gestational ‘potassium switch’. **(b)** Intracellular calcium is regulated by SR up-take (SERCA) and release (RyR/IP_3_R). **(c)** Cells are electrically coupled via gap junctions between excitable cells (*κ*) and excitable with passive (*p*_*gp*_).

### B. Dimensional Analysis

To facilitate efficient numerical integration and identify the dominant physiological timescales, we non-dimensionalised the system. Detailed derivations are provided in **Appendix J**. This process yielded a set of dimensionless groups (denoted as Π) that mathematically isolate the competing biophysical mechanisms governing the transition to labour. These groups, defined in Table 1, bridge the fast intracellular channel kinetics (Π_*τ*_) to the slow tissue-wide wave propagation. Crucially, this formulation explicitly separates two topologically and functionally distinct intercellular pathways: the synchronising USMC-USMC gap junction network (Π_*κ*_) and the stabilising, hyperpolarising current sink between USMCs and interstitial cells (**the passive cell network**) (Π_*p*_).

**Table 1.** Key Dimensionless Groups governing the USMC-FTU model. Definitions correspond to derivations in Appendix A.

| Group | Definition | Physiological Interpretation |
| --- | --- | --- |
| **Gating Speed** ($\Pi_\tau$) | $\frac{\tau_0}{\tau_\chi}$ | **Timescale Separation:** Indicates if a channel gate is fast ($\gg 1$) or slow ($\ll 1$) relative to the system characteristic time ($\tau_0 = 1.0$ s). |
| **Excitability Ratio** | $\frac{g_{CaT}}{g_{Kvm}}$ | **Pacemaker Potential:** The balance between the destabilising pacemaker drive ($I_{CaT}$) and the stabilising repolarisation reserve ($I_{Kvm}$). |
| **Quiescence Clamp** ($\Pi_p$) | $\frac{p_{gp}\tau_0}{C_m}$ | **Stabilising Sink:** Represents the strength of the current sink provided by the passive cell network; this must decrease to permit spontaneous firing. |
| **Coupling Strength** ($\Pi_\kappa$) | $\frac{\kappa\tau_0}{C_m}$ | **Synchronisation:** The efficiency of charge transfer between active cells, governing the spatial propagation of the action potential. |

### C. Physiological Validation Criteria (Parameter Selection)

To account for the intrinsic cellular heterogeneity of the myometrium, a single set of parameters is insufficient to capture the full spectrum of cellular phenotypes present in the tissue. Instead, we utilised an ensemble approach to identify a population of physiologically valid baseline models capable of undergoing the full gestational transition.

From an initial ensemble of 100,000 parameter sets (generated by varying conductances by *±*100% around the baseline), we filtered cells based on three gestational milestones:

1. **Quiescence (Evaluated at representative day 160; 22 weeks, 6 days):** Cells must remain stable and non-bursting under mid-gestation conditions (Second Trimester).
2. **Excitability (Evaluated at representative day 250; 35 weeks, 5 days):** Cells must demonstrate the capacity for action potential generation when stimulated (Third Trimester).
3. **Spontaneous Bursting (Evaluated at representative day 275; 39 weeks, 2 days):** Under late-term conditions (reduced *g*_*Kvm*_), cells must exhibit spontaneous, repetitive bursting.

Only 34 baseline parameter sets survived this rigorous filtering process. A parameter set was deemed successful only if a continuous simulation across gestation, driven dynamically by the gestational transition functions (defined in Section E), sequentially satisfied all three milestones.

### D. Functional Tissue Unit Architecture

Tissue-level interactions were simulated using a 2D network of coupled cells, an FTU, was defined as a 7 × 7 lattice (*N* = 49 cells). While macroscopically small, this dimension was explicitly calculated to be the minimal functional domain that provides a structurally dominant core (cells with complete neighbour coupling) to prevent artificial boundary-driven excitability. This specific scale ensures the emergent burst frequency is governed by the intrinsic pacemaker distribution rather than edge effects, while remaining computationally tractable for solving 588 stiff ODEs across 100,000 ensemble simulations (see Appendix C for formal mathematical justification). Because the primary driver of rhythmic frequency in this regime is the cellular excitability ratio rather than spatial wave-front collision, simulating larger lattices would increase computational cost without fundamentally altering the baseline pacemaking frequency established by this minimal unit.

The lattice comprises two cell types:

1. **Active USMCs:** Cells capable of contraction and potential pacemaker activity.
2. **Passive Cells:** Representing electrical loads or interstitial cells Means et al. (2025); Duquette et al. (2005).

Intercellular communication within this architecture occurs via two functionally distinct pathways, which are explicitly separated by our mathematical formulation. The dimensionless group Π_*κ*_ governs the synchronising, spatially heterogeneous gap junction network between adjacent USMCs (*κ*) Tabb et al. (1992). In contrast, Π_*p*_ governs the stabilising, hyperpolarising current sink between USMCs and their coupled passive cells (*p*_*gp*_).

### E. Gestational Remodelling and Evolving Heterogeneity

A critical feature of this model is its ability to simulate the profound electrophysiological transformation of the myometrium from quiescence to term. Rather than modelling a single, monolithic transition, our model employs three distinct, phenomenological sigmoidal functions to represent the different maturation timelines of key physiological processes. Crucially, these continuous functions act as time-dependent multipliers applied to the 34 validated baseline parameter sets (seeds), dynamically evolving their properties across the simulated gestational timeline. Each function, *f* (*t*_*gest*_), follows the general form:

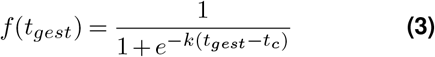

where *t*_*gest*_ is the gestational day, *t*_*c*_ is the midpoint of the transition, and *k* controls its steepness. The three specific transition functions are:

1. **Pacemaker Potential (***f*_*pacemaker*_): (*t*_*c*_ = 160 days; Second Trimester, *k* = 0.045). Governs the gradual development of the pacemaker sub-population, increasing the proportion of cells with high *g*_*CaT*_ Zhu et al. (2023) and reduced *g*_*Kvm*_ during the second trimester.
2. **General Excitability (***f*_*excitability*_): (Inflection point *t*_*c*_ = 220 days; Early Third Trimester, *k* = 0.03). Controls the bulk tissue properties—reaching the validated excitable state by Day 250—including the upregulation of L-type calcium channels (*g*_*CaL*_) Wray and Arrowsmith (2021) and the critical decrease in USMC-passive cell coupling (*p*_*gp*_).
3. **Coordination (***f*_*coord*_): (*t*_*c*_ = 275 days; Late Third Trimester/Term, *k* = 0.15). Models the rapid, sharp upregulation of Connexin-43 gap junctions (*κ*) immediately prior to term. This explicitly enhances USMC-USMC synchronisation, representing a distinct intercellular pathway from the USMC-passive cell coupling (*p*_*gp*_) that is concurrently downregulated to remove the quiescence clamp Garfield et al. (1988); Tabb et al. (1992).

These functions modulate both the bulk tissue properties and the emergence of the pacemaker sub-population, as illustrated in Figure 2, and drive the time-dependent changes in model parameters summarised in Table 2.

**Table 2.**
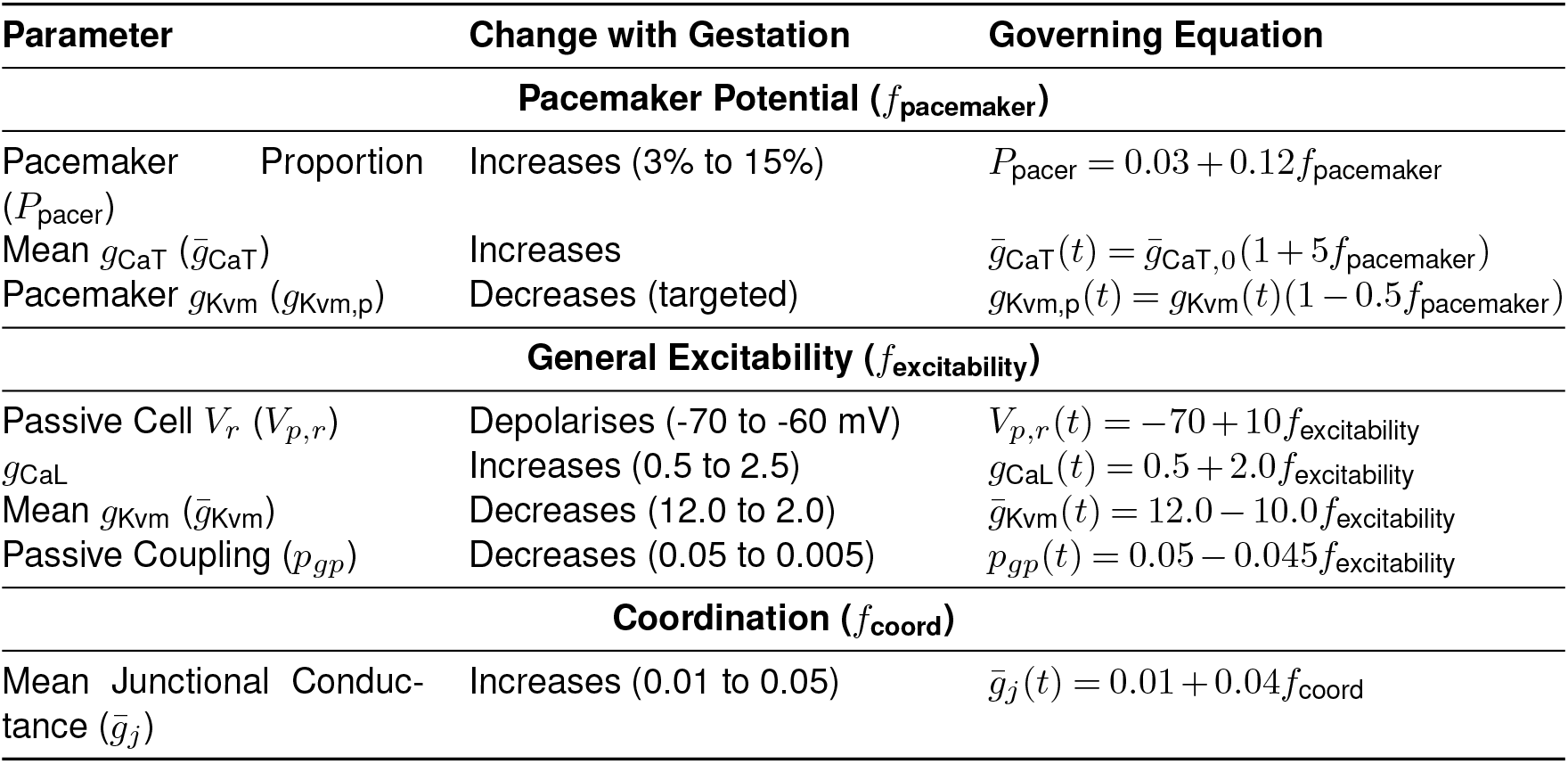
Summary of Gestational Remodelling of Model Parameters via Transition Functions.

**Figure 2.**
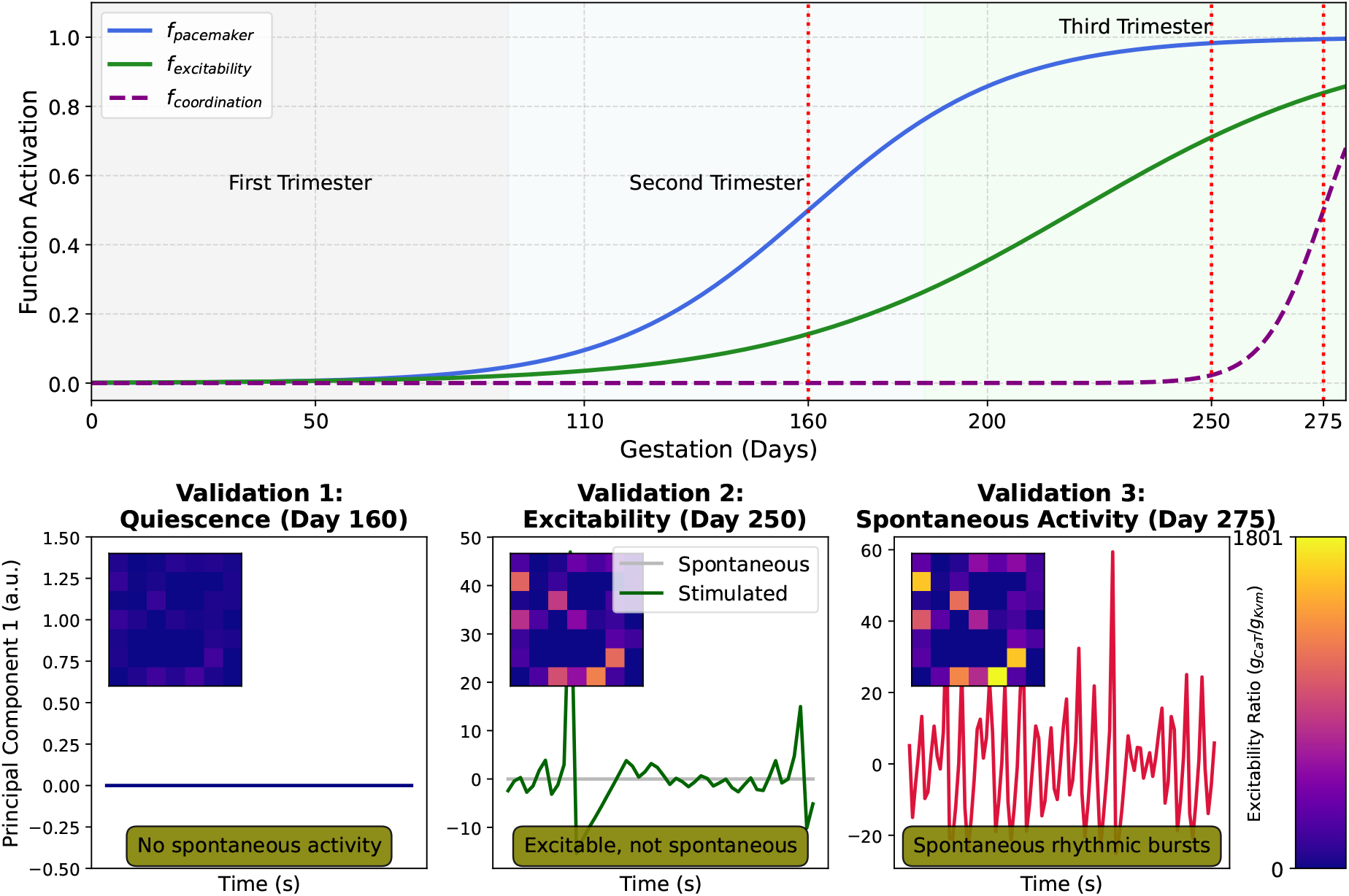
Methodological Framework and Validation of the Gestational Myometrial Model. **(Top Panel)** A gestational timeline showing the three phenomenological functions (*f*_pacemaker_, *f*_excitability_, *f*_coordination_) that govern remodelling. Red dotted vertical lines indicate the validation days (*t*_*c*_ = Second-, Early-Third-, and Late-Third-Trimester). **(Bottom Panels)** The model’s emergent electrical behaviour (PC1) at each validation point: (left) Quiescence (stable and non-excitable); (middle) Excitability (capable of induced propagation but lacking spontaneous bursting); (right) Spontaneous Bursting. **(Insets)** Visualisation of the cellular excitability ratio (*g*_*CaT*_ */g*_*Kvm*_) on the 7 × 7 grid, illustrating the dynamic emergence of the pacemaker sub-population. This ratio progressively shifts from a repolarisation-dominated baseline to a highly excitable, depolarisation-dominated state at term (see Figure 3B for the quantitative distribution of these values across the validated ensemble).

The progressive depolarisation of the passive cell resting potential (*V*_*p,r*_) from a highly hyperpolarised mid-gestation state of − 70 mV to a term state of −60 mV captures the broad electrophysiological shift of the myometrial cellular ecosystem. This late-gestation target of −60 mV is specifically chosen to fall within the experimental range (−58*±*7 mV) reported in patch-clamp recordings of uterine interstitial cells Duquette et al. (2005). Furthermore, while the resting potential of uterine fibroblasts varies widely, the majority reside in an even more depolarised state above −25 mV Xu et al. (2015). Physiologically, this shift to −60 mV does not actively drive excitation; rather, it effectively weakens the quiescence clamp. By reducing the electrotonic gradient between the passive sink and the USMC, it drains less depolarising current from the active cells, thereby permitting the intrinsic pacemaking mechanism to successfully initiate a burst.

### F. Simulation Protocols

#### Pacemaker Selection

We tested the hypothesis of heterogeneity-driven pacemaking by identifying the USMC within the lattice possessing the highest intrinsic T-type calcium conductance (*g*_*CaT*_) Lee et al. (2009) and selectively reducing its potassium conductance (*g*_*Kvm*_), mimicking the potassium switch Green-wood and Tribe (2014).

#### Inflammation Experiment

To simulate pathology, we introduced a perturbation to the ion channel parameters consistent with cytokine-induced remodelling (specifically the modification of potassium channel conductance Khan et al. (1993) and the upregulation of Connexin-43 associated with inflammatory cytokines Sivarajasingam et al. (2016)), assessing whether this shift was sufficient to trigger preterm bursting events.

### G. Data Analysis and Signal Processing

To quantify tissue-level synchrony from the heterogeneous activity of the *N* = 49 cells, we performed principal component analysis (PCA) on the ensemble voltage matrix **V**(*t*) ∈ ℝ^*N* ×*T*^ Raine et al. (2015). The first principal component (PC1) was extracted as the effective waveform, representing the dominant mode of coordinated tissue activity. Burst frequency was subsequently calculated from the envelope of the PC1 signal, derived using the Hilbert transform, to robustly identify burst onset and termination even in complex waveform shapes Boashash (1992).

## Results

### Emergence of Tissue-Level Function from a Selected Pacemaker Population

Out of the 100,000 seeds simulated, only 34 successfully met the three-stage physiological milestones (Quiescence → Excitability → Bursting). Figure 3A shows a representative time-series from one of these successful seeds (Seed 2709). The effective waveform (PC1) at Day 275 demonstrates robust, high-amplitude rhythmic bursting, whereas the waveforms for Day 160 and Day 250 (insets) show quiescence and non-bursting excitability, respectively. Figure 3B illustrates the distribution of these 34 successful seeds within the parameter space defined by the excitability ratio (*g*_*CaT*_ */g*_*Kvm*_) versus the pacemaker population size. The successful seeds (green stars) cluster in a specific region distinct from the failing seeds (background density), characterised by a high intrinsic *g*_*CaT*_ in the pacemaker sub-population.

**Figure 3.**
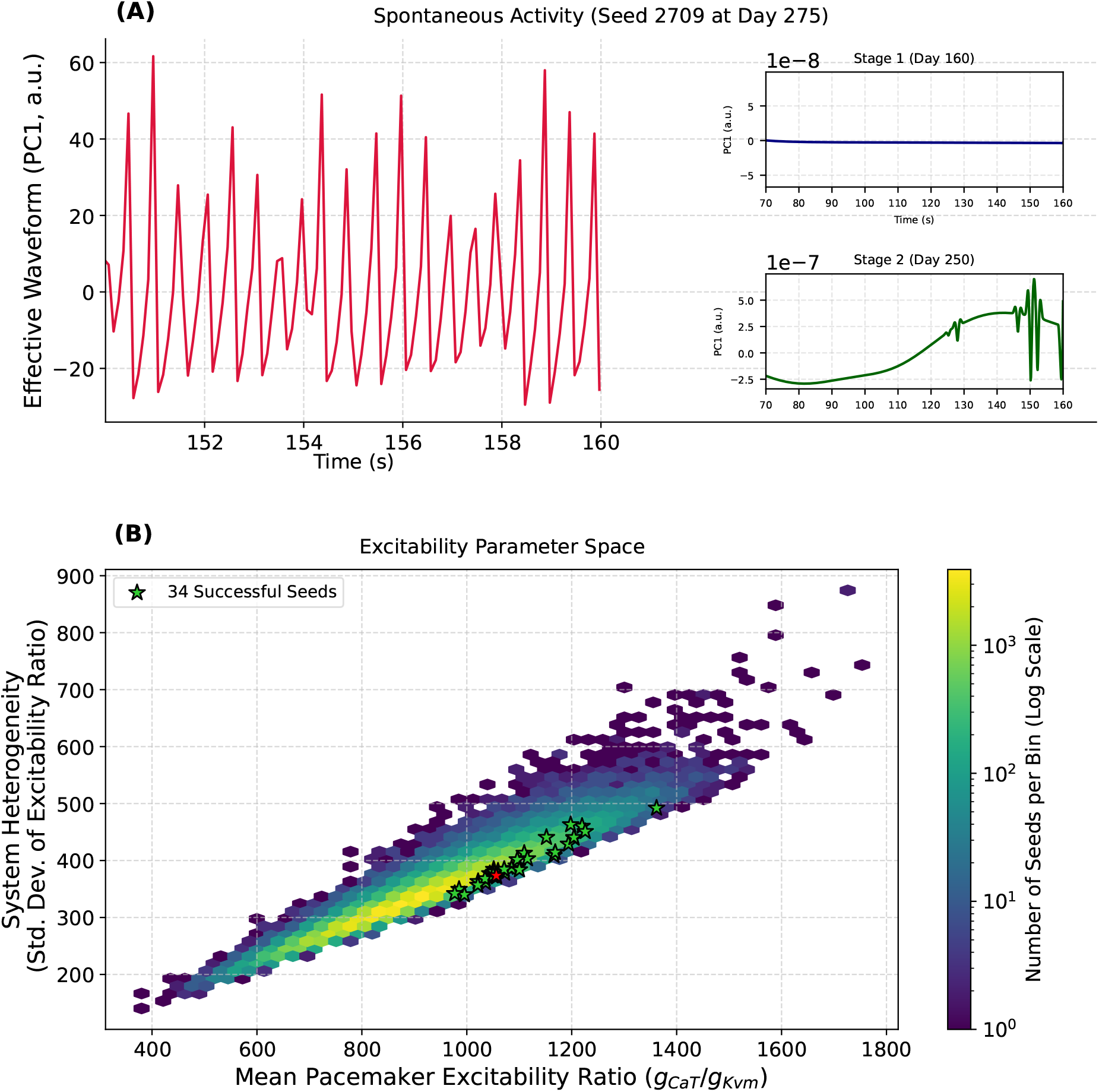
**(A)** Initiation of spontaneous activity from a successful simulation (Seed 2709, Day 275); only the last 10 seconds of the 160-second simulation are shown to highlight the burst morphology. Across its full duration, this seed yields a physiological frequency of *≈* 2.8 contractions per minute, matching clinical targets for active labour (comprehensive ensemble statistics are detailed in Figure 4). The insets show the effective waveform (PC1) for Day 160 (Quiescent) and Day 250 (Excitable but non-bursting). **(B)** The parameter space of pacemaker cell population excitability versus the heterogeneity of the excitability ratio for all 100,000 simulated seeds. The 34 successful seeds (green stars) cluster in a region of high pacemaker excitability. Notably, this specific heterogeneity profile only yields spontaneous bursting because the bulk USMC-passive coupling (*p*_*gp*_) has concurrently decreased at term to release the quiescence clamp.

To test the sensitivity of this emergent behaviour to the specific spatial arrangement of connections, we performed a robustness check.

For all 34 successful seeds, the stochastic cell-cell coupling weights (*κ*_*i,base*_) were randomly reshuffled while maintaining the same statistical distribution. Specifically, these weights were sampled from a positive half-normal distribution (*µ* = 0, *σ* = 1), ensuring intercellular coupling was heterogeneously distributed but strictly positive (non-zero) across the tissue. In 100% of these reshuffled cases, the tissue retained its ability to generate spontaneous, propagating bursts at Day 275, satisfying the physiological milestones.

### I.Analysis of Validated Simulation Ensemble

Quantitative analysis of the 34 successful seeds revealed a consistent electrophysiological profile at term (Day 275). As shown in Figure 4A, the mean burst frequency was 0.047*±*0.01 Hz (≈2.8 contractions per minute). During these bursts, the mean peak intracellular calcium concentration reached 0.145*±*0.012 µM. In terms of tissue architecture, these successful configurations were characterised by a mean pacemaker-to-follower excitability ratio (*g*_*CaT*_ */g*_*Kvm*_) of 1100*±*97.

**Figure 4.**
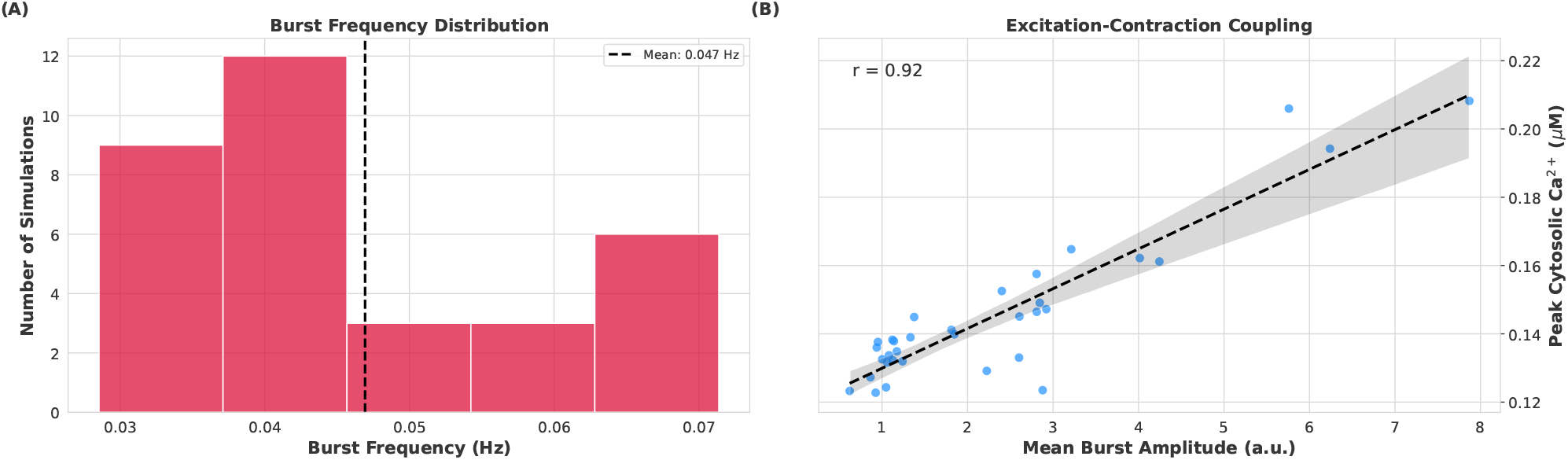
Analysis of Emergent Burst Dynamics at Day 275. **(A) Burst Frequency Distribution:** The mean frequency (0.047 Hz) falls within the target clinical range for active labour (2–3 contractions per minute, corresponding to roughly 0.033–0.050 Hz), which spans the bulk of the simulated distribution. **(B) Excitation-Contraction Coupling:** A scatter plot showing the correlation (*r* = 0.92) between the mean amplitude of electrical bursts (PCA mode) and the peak intracellular Ca^2+^ concentration.

We further examined the relationship between electrical activity and calcium dynamics. As expected from standard excitation-contraction coupling, Figure 4B demonstrates a strong positive correlation (*r* = 0.92) between the amplitude of the electrical bursts (PC1 amplitude) and the peak intracellular Ca^2+^ concentration. However, the primary functional relevance of including SR calcium handling in this model lies in its temporal, rather than amplitude-generating, dynamics. While some early animal studies have suggested SR calcium release may be dispensable for the bare initiation of spontaneous activity (e.g., in rats Shmigol et al. (2001)), the intracellular calcium clearance rate governed by SR uptake (Π_*flux*_) is essential in our human FTU formulation. It critically governs the refractory period and drives burst termination, ensuring the tissue reproduces the distinct, separated rhythmic bursts of labour rather than entering continuous electrical tetanus. Crucially, achieving these quantitative milestones represents more than just model validation; it provides computational evidence for a major physiological hypothesis. These results demonstrate that an active, intrinsic USMC mechanism—driven by the *potassium switch* and functional decoupling from the interstitial network—is entirely sufficient to drive term labour contractions. Furthermore, the shuffling experiments prove that this rhythmicity is an emergent property of the tissue’s statistical distribution of electrical properties rather than its specific spatial micro-architecture. Consequently, the model reveals that the myometrium does not require a fixed, vulnerable anatomical pacemaker to function, but could instead operate as a highly redundant, self-organising syncytium.

### J. Simulating an Inflammation-Induced Preterm Labour Phenotype

To assess the model’s response to pathological parameter shifts, we simulated an inflammation scenario at Day 180 (normally quiescent). Starting from a control state where the tissue was quiescent (Figure 5A), we introduced two targeted perturbations to mimic the acute effects of pro-inflammatory cytokines: (1) a 50% reduction in the repolarising BK channel conductance (*g*_*bka*_), reflecting the well-documented cytokine-mediated suppression of potassium channel expression Khan et al. (1993), and (2) a premature increase in mean gap junction conductance 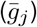 to term levels (0.05 µS), simulating inflammation-driven Connexin-43 upregulation Sivarajasingam et al. (2016). As shown in Figure 5B, these specific molecular-level changes were sufficient to transition the tissue from quiescence to a state of high-amplitude, spontaneous bursting, effectively reproducing a preterm labour phenotype *in silico*.

**Figure 5.**
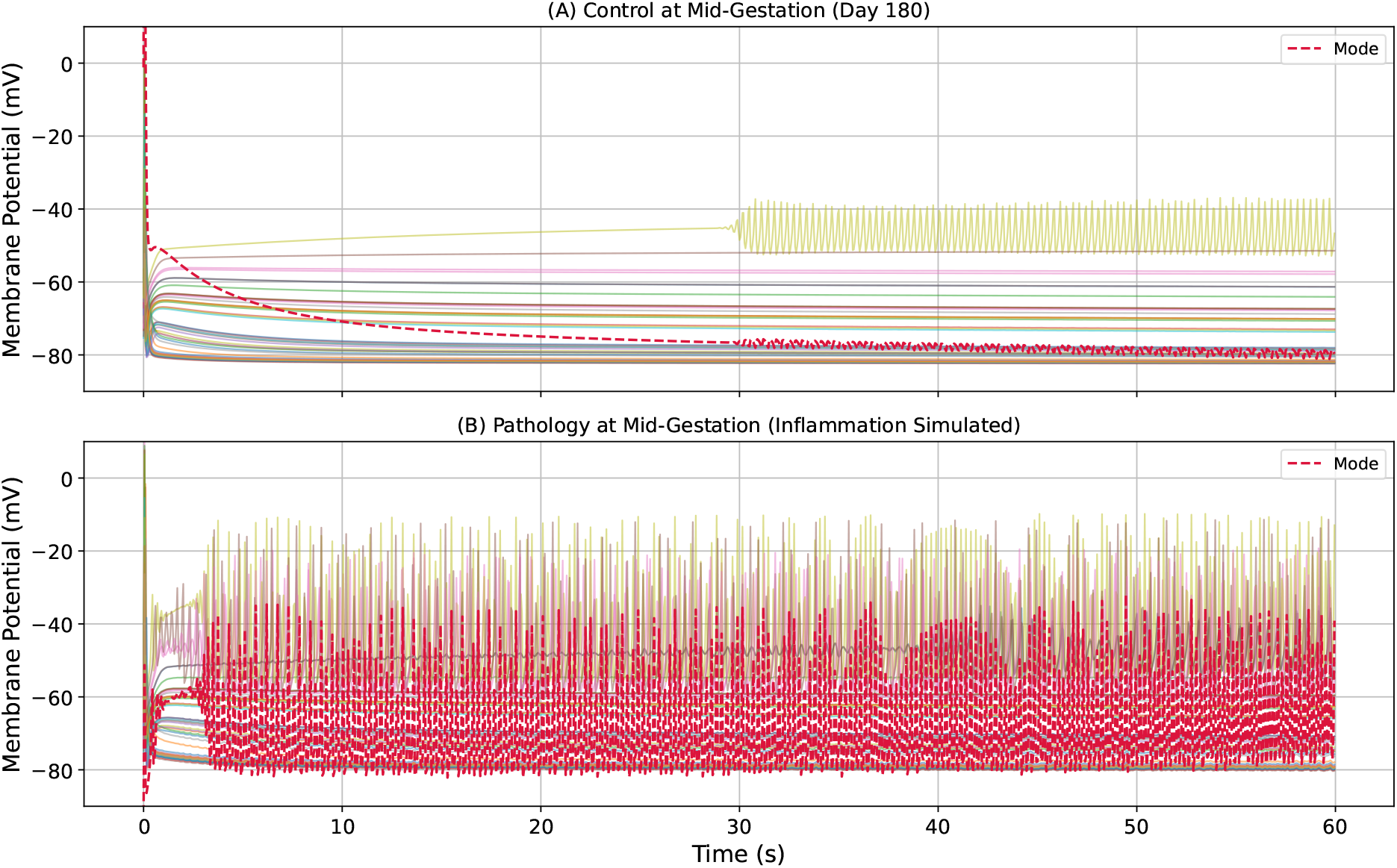
Simulating an Inflammation-Induced Preterm Labour Phenotype. **(A) Control (Day 180):** The tissue is quiescent. **(B) Inflammation (Day 180):** Following the reduction of *K*^+^ conductance and increase in gap junction coupling, the tissue transitions to spontaneous, rhythmic bursting. In both panels, the thin, multi-coloured solid lines represent the individual membrane potential traces of the 49 simulated cells comprising the FTU. The bold crimson dashed line indicates the dominant tissue-level waveform extracted via Principal Component Analysis (PC1, denoted as ‘Mode’ in the legend).

## Discussion

This study presents a multi-scale computational framework to investigate the transition of the human myometrium from gestational quiescence to the synchronised contractility of labour. By integrating biophysically detailed cellular models into a functional tissue unit, we demonstrated that a heterogeneity-driven pace-maker selection mechanism is sufficient to generate robust, spontaneous rhythmicity. Our results show that the emergence of a specific sub-population of USMCs—defined by high intrinsic *g*_*CaT*_ and reduced *g*_*Kvm*_—can entrain the bulk tissue, producing coordinated bursts at a frequency of 0.047 Hz (≈2.8 min^*−*1^). This frequency closely aligns with clinical observations of active labour Devedeux et al. (1993); Buhimschi (2009), validating the model’s temporal dynamics against physiological benchmarks. Crucially, this emergent function was achieved without imposing a predefined anatomical pacemaker, corroborating the hypothesis that the uterine pacemaker is a dynamic, functional entity rather than a fixed structural one, capable of shifting location as local excitability gradients evolve.

A central question in uterine physiology is the identity of the primary driver of contractions: is it an extrinsic network of interstitial cells or an intrinsic mechanism within the smooth muscle itself? Our model explicitly investigated the latter by configuring the passive cell network strictly as a hyperpolarising electrotonic load (anchored at a telocyte-like −60 mV, isolating this from potentially stimulatory fibroblasts near −15 mV). Under these constraints, demonstrating that USMC re-modelling alone can overcome this load to sustain coordinated bursting provides robust evidence for the intrinsic sufficiency of the myometrium. Critically, this mechanism relies on the physiological hypothesis that bulk USMC-passive coupling (*p*_*gp*_) is down-regulated independently of the concurrent term up-regulation in homotypic USMC-USMC gap junctions (*κ*). This divergent regulation is biologically grounded: while USMC-USMC connections are primarily homotypic (Cx43), USMC-interstitial connections form heterotypic junctions (e.g., Cx43/Cx45). Biophysical and computational studies of the myometrium demonstrate that such heterotypic gap junctions exhibit distinct, typically lower unitary conductances and asymmetric voltage-gating, allowing them to act independently as negative modulators of excitability Martinez et al. (2002); Sheldon et al. (2014). Consequently, our computational results highlight this disparate, targeted regulation of heterotypic versus homotypic connexin conductances as a critical avenue for future experimental validation, proposing it as a potent functional switch not only in parturition but potentially across other mixed-cell physiological syncytia. This suggests that while interstitial cells likely play a significant modulatory or distinct pacemaking role, they are not a strict prerequisite for rhythmicity if the USMC population undergoes sufficient ion channel remodelling. We propose that the uterus likely operates with a physiological fail-safe mechanism: the active potassium switch and heterogeneity-driven selection provide a robust intrinsic drive that ensures parturition can proceed even if the passive interstitial network is compromised or functionally decoupled. This redundancy is evolutionarily advantageous for an organ where failure of function is critical.

The robustness of this intrinsic drive was further evidenced by the shuffling experiments, which challenged the model’s reliance on specific micro-architecture. The finding that spontaneous bursting persisted despite the random spatial permutation of the USMC-USMC gap junction weights (*κ*_*i,base*_) offers a critical mechanistic insight. Importantly, this shuffling applied exclusively to the USMC syncytium, while the passive cell network properties (*p*_*gp*_ and *V*_*p,r*_) were maintained as uniform, bulk electrotonic loads across the FTU. This demonstrates that at the micro-scale, tissue-level initiation is an emergent property of the statistical distribution of cellular phenotypes, rather than a specific micro-structural connectivity pattern. While macro-scale spatial gradients in passive coupling are undoubtedly critical for orienting propagating waves across the whole organ (as demonstrated in previous large-scale lattice models Means et al. (2025)), our FTU results imply that the local *initiation* of a burst does not require a precise, hard-wired cellular circuit. Such a rigid requirement would be fragile and prone to failure during the immense physical distension of pregnancy. Instead, local initiation relies on a global upregulation of mean connectivity and excitable ion channel density. As long as the mean homotypic coupling is sufficient to allow the pacemaker sub-population to overcome this strictly defined hyperpolarising sink, the precise local spatial arrangement of these connections is secondary for local burst initiation. Our analysis of the dimensionless parameter space reveals that this onset of labour may be described as a threshold phenomenon. As explicitly visualised in the parameter space of Figure 3B, successful transition to bursting requires the pacemaker sub-population to cross a specific excitability manifold (empirically bounded at a mean *g*_*CaT*_ */g*_*Kvm*_ ratio *>* 990) where its intrinsic depolarising drive (Π_*CaT*_) strictly exceeds the stabilising influence of the bulk quiescence clamp (Π_*p*_). The specific mechanistic roles of these competing dimensionless groups are summarised in Table 3. Consequently, if *p*_*gp*_ remains high at term, Π_*p*_ mathematically dominates; the passive sink drains the nascent electrical activity, rendering the cellular excitability ratio irrelevant and locking the tissue in quiescence.

**Table 3.** Dimensionless Groups and Their Roles in System Stability.

| Group | Role in Stability | Mechanism of Action |
| --- | --- | --- |
| Excitability Ratio<br>($\Pi_{CaT}/\Pi_{Kvm}$) | **Pacemaker Drive** | Balances spontaneous depolarisation ($\Pi_{CaT}$) against the repolarising restoring force ($\Pi_{Kvm}$); high ratios overcome the stable fixed point (quiescence) and drive the system towards a limit cycle (oscillatory) regime. |
| $\Pi_p$ | **Stabiliser (Clamp)** | Acts as a current sink; couples the USMC to the stable resting potential of the passive network, suppressing small fluctuations. |
| $\Pi_{\kappa}$ | **Coupling/Synchroniser** | Facilitates charge diffusion; allows local excitations to recruit neighbours, overcoming the local electrotonic load. |
| $\Pi_{flux}$ | **Refractory Controller** | Governs $Ca^{2+}$ clearance rate; determines the duration of the refractory period and burst termination. |

By maintaining this uniform hyperpolarising load, our formulation provides an essential, micro-scale complement to previous macro-scale lattice models. For instance, our earlier work using abstract FitzHugh-Nagumo models demonstrated that macroscopic spatial heterogeneities in *p*_*gp*_ and a depolarised passive network are vital for *orienting* and steering tissue-level waves Means et al. (2025). In stark contrast, to rigorously prove *intrinsic* initiation, our biophysical FTU deliberately stripped away these extrinsic spatial gradients and stimulatory voltages. By demonstrating that rhythmic bursts can ignite locally even against a uniform, hyperpolarised sink, we establish the baseline cellular engine. At this micro-scale, the temporal functional uncoupling of USMCs from the passive interstitial sink (the global gestational decrease in bulk *p*_*gp*_) is just as critical as the upregulation of homotypic USMC-USMC coupling (*κ* ↑) Sheldon et al. (2013); Xu et al. (2015). Thus, the potassium switch (*g*_*Kvm*_ ↓) and interstitial decoupling (*p*_*gp*_ ↓) must occur in concert to permit the phase transition to active labour.

While this functional tissue unit model captures the essential dynamics of the transition to labour, several limitations remain that delineate the scope for future work. First, the current 2D lattice structure simplifies the complex, anisotropic 3D fibre architecture of the real myometrium Pullan et al. (2010). Future iterations will embed this formulation within anatomically realistic finite element models to investigate how fibre orientation influences wave propagation vectors. Second, the model employs a generic description of Sarcoplasmic Reticulum calcium release. The emergent peak cytosolic calcium during bursts (0.145*±* 0.012 µM) was sufficient to trigger standard excitation-contraction coupling cascades; however, it lies below the higher physiological estimates (200–800 nM) typical of robust contractions Wray et al. (2003). This discrepancy arises because the current FTU lacks explicit IP_3_-receptor mediated calcium-induced calcium release (CICR) Arrow-smith and Wray (2014); Kimura et al. (1996). While integrating CICR would serve as a critical non-linear amplifier to match the true force generation magnitude required for the expulsive phase of labour, its absence does not invalidate the primary temporal findings. The underlying electrophysiological pacemaking frequency (0.047 Hz) is fundamentally governed by the voltage-gated clock and repolarisation reserve, meaning an added calcium amplification factor would scale the transient amplitude without disrupting the fundamental rhythmicity established here. Finally, we simplified the inflammatory pathway into a phenomenological parameter shift. While this successfully reproduced a preterm phenotype, a more mechanistic approach integrating specific cytokine signalling cascades (e.g., IL-1*β*, TNF-*α*) would allow for more predictive *in silico* drug testing Sivarajasingam et al. (2016); Mor et al. (2017).

In conclusion, this study establishes a quantitative framework for the myometrial functional tissue unit. By validating the sufficiency of heterogeneity-driven pace-making, we offer a reconciled view of uterine activation where intrinsic USMC remodelling provides a robust, redundant driver of parturition. This platform serves as a foundational module for future whole-organ simulations within the Physiome Project Hunter and Borg (2003), bridging the gap between molecular channelopathies and clinical dystocia.

## Ethics

This study is entirely computational and did not involve new experiments on human or animal subjects; therefore, ethical approval was not required.

## Data accessibility

The numerical implementation of mathematical model and simulation code used to generate the results in this study are available on git hub at URL https://github.com/Jagirhussan/uterus-ftu

## Authors’ contributions

J.R.H. conceptualized the study, developed the mathematical model, performed the simulations, and wrote the initial draft of the manuscript. S.A.M. codeveloped the mathematical model, contributed to the methodology and data analysis. P.J.H. and A.R.C. provided supervision, funding acquisition, and critical interpretation of the results. All authors contributed to the editing and critical revision of the manuscript, and all gave final approval for publication.

## Acknowledgements

The authors wish to acknowledge the use of New Zealand eScience Infrastructure (NeSI) high-performance computing facilities, consulting support and/or training services as part of this research. New Zealand’s national facilities are provided by NeSI and funded jointly by NeSI’s collaborator institutions and through the Ministry of Business, Innovation & Employment’s Research Infrastructure programme. URL https://www.nesi.org.nz.

## Appendix: Mathematical Derivations and Model Details

### A. The Dimensional Functional Tissue Unit Model

The model consists of a coupled system of ordinary differential equations (ODEs) describing the active Uterine Smooth Muscle Cell (USMC) and the passive interstitial cell.

#### State Variables

The system is defined by:

- **Membrane Potentials:** *V*_*m*_ (Active USMC), *V*_*p*_ (Passive Cell).
- **Calcium Dynamics:** *C*_*i*_ (Cytosolic Ca^2+^), *C*_*SR*_ (Sarcoplasmic Reticulum Ca^2+^).
- **Gating Variables (***χ*): 15 state variables governing ion channel kinetics (*m, h, d, f*_1_, *f*_2_, *b, g, y, q, r*_1_, *r*_2_, *p, k*_1_, *k*_2_, *s, x*_*a*_, *x*_*ab*_, *c*).

#### Model Order Reduction

The full biophysical formulation of the active USMC originally comprises 18 state variables. However, to reduce extreme computational stiffness during tissue-level simulations, we apply the quasi-steady-state approximation (QSSA) to the fastest gating variables prior to numerical integration. As formally justified by the non-dimensionalisation process (see **Derivation 1**B), variables operating on a much faster timescale (Π_*τ*_ ≫ 1) than macroscopic tissue dynamics are computed algebraically via their steady-state voltage-dependent functions (*x ≈ x*_*∞*_(*V*_*m*_)).

#### Variables approximated via QSSA

- *m* (*I*_*Na*_ activation)
- *d, f*_1_ (*I*_*CaL*_ activation and fast inactivation)
- *b, g* (*I*_*CaT*_ activation and inactivation)

#### Post-QSSA Dynamically Integrated Variables

Applying this timescale reduction leaves exactly 12 ordinary differential equations (ODEs) per cell unit to be solved computationally. For the complete 7 × 7 FTU lattice (*N* = 49), this yields precisely 12 × 49 = 588 stiff ODEs. The 12 dynamically integrated state variables per unit are:

- **Membrane Potentials:** *V*_*m*_ (Active USMC), *V*_*p*_ (Passive Cell).
- **Calcium Dynamics:** *C*_*i*_ (Cytosolic Ca^2+^), *C*_*SR*_ (Sarcoplasmic Reticulum Ca^2+^).
- **Slow Gating Variables:** *h* (*I*_*Na*_ inactivation), *q, r*_1_, *r*_2_ (*I*_*K*1_ activation and inactivation), *f*_2_ (*I*_*CaL*_ slow inactivation), *c* (*I*_*Cl*(*Ca*)_ gating), *r*_*rel*_ (RyR gating), and *s*_*serca*_ (SERCA gating).

##### 1. Membrane Potential Equations

The time evolution of the transmembrane voltage is governed by the conservation of charge. **Active Cell (***V*_*m*_):

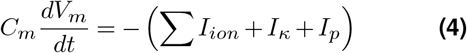

where *I*_*κ*_ is the USMC-USMC gap junction current and *I*_*p*_ is the USMC-Passive coupling current.

###### Passive Cell Equations (Derivation)

The passive cell is modelled as an equivalent RC circuit lacking voltage-gated ion channels. Its membrane potential (*V*_*p*_) is governed by the balance of two currents: a passive leakage current (*I*_*leak*_) and the intercellular coupling current (*I*_*p*_) from the active cell.

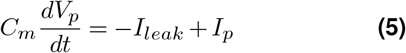

Substituting the Ohmic formulations *I*_*leak*_ = *g*_*leak*_(*V*_*p*_ *− V*_*p,r*_) and *I*_*p*_ = *p*_*gp*_(*V*_*m*_ *− V*_*p*_):

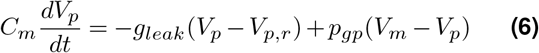

Dividing by the capacitance *C*_*m*_, we obtain the governing rate equation:

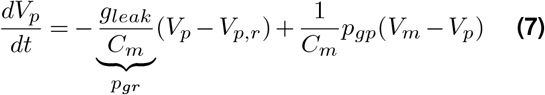

Here, the parameter *p*_*gr*_ = *g*_*leak*_*/C*_*m*_ represents the inverse membrane time constant 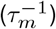, physically interpreted as the relaxation rate of the potential towards its resting state *V*_*p,r*_ in the absence of coupling.

##### 2. Ionic Current Formulations

The total ionic current Σ*I*_*ion*_ comprises the following components, based on the Tong *et al*. (2011) formulation:

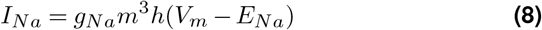

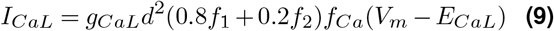

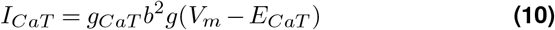

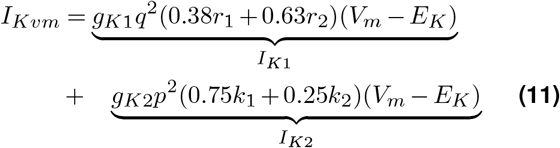

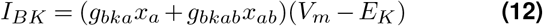

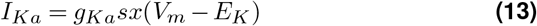

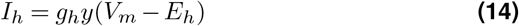

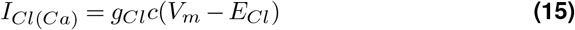

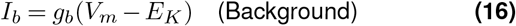

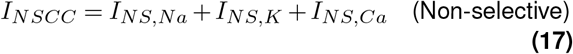

*Note: I*_*CaL*_ includes calcium-dependent inactivation via the factor 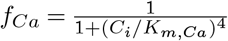. Equations (9–11) represent the primary currents modulated during gestational remodelling.

##### 3. Calcium Dynamics

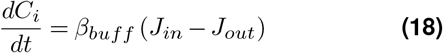

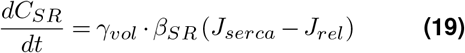

###### Flux Definitions

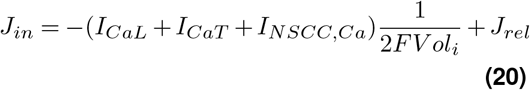

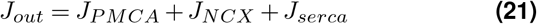

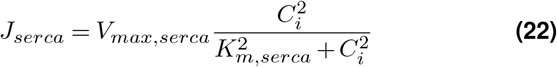

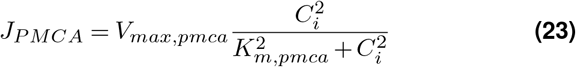

#### B. Non-Dimensionalisation Analysis

To identify the dominant stiffness in the system and enable efficient numerical integration, we scaled the dimensional equations using the following characteristic scales:

- **Time Scale** (*τ*_0_): 1.0 s (Representative of the slow tissue-level contraction timescale).
- **Voltage Scale** (*V*_0_): *RT/F ≈* 26.7 mV (Thermal voltage).
- **Concentration Scale** (*C*_0_): 0.1 µM (Order of magnitude for basal cytosolic calcium).

This transformation yields the dimensionless variables: 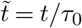, 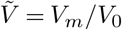, and 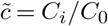.

The dimensional system is highly stiff, typically requiring implicit solvers. However, the non-dimensionalisation process explicitly separates the fast (Π_*τ*_ ≫ 1) and slow (Π_*τ*_ ≪ 1) timescales. This separation guided our QSSA reduction, allowing us to utilise an explicit Runge–Kutta (RK45) solver with adaptive stepping, significantly reducing computation time.

##### Derivation 1: Timescale Separation (Π_τ_)

For any gating variable *χ*, the kinetics are described by 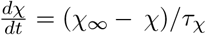. Substituting the dimensionless time 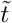:

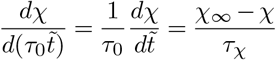

Multiplying by *τ*_0_ isolates the dimensionless group governing stiffness:

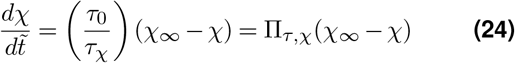

##### Interpretation: Π_*τ,χ*_ represents the ratio of the macroscopic system timescale to the intrinsic channel timescale

- If Π_*τ*_ *≫* 1 (e.g., *I*_*Na*_ activation), the process is fast and can be approximated by QSSA.
- If Π_*τ*_ ≪ 1 (e.g., *I*_*K*_ inactivation), the process is slow and rate-limiting.

##### Derivation 2: Relative Conductance (Π_ion_)

We scale the voltage conservation equation:

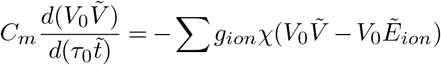

where 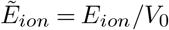 is the dimensionless Nernst potential. Dividing through by the coefficient term 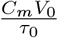:

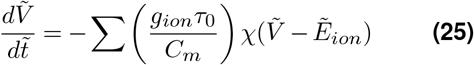

This isolates the dimensionless conductance group:

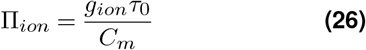

This group quantifies the strength of a specific ion channel relative to the membrane capacitance. It applies analogously to the intercellular coupling terms:

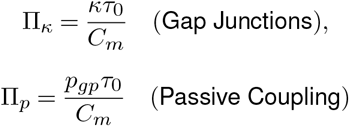

##### Derivation 3: Calcium Transport (Π_flux_)

For pump fluxes (e.g., SERCA or PMCA) following Michaelis–Menten kinetics, the scaling of the mass balance equation 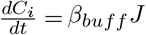 proceeds as:

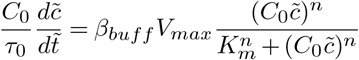

Rearranging yields:

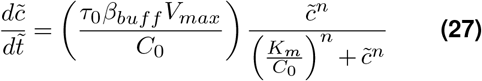

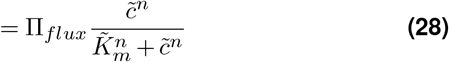

Here, Π_*flux*_ represents the dimensionless capacity of the pump to alter cytosolic calcium levels within the characteristic timescale *τ*_0_, and 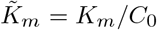 is the dimensionless Michaelis constant.

#### C. Justification for Lattice Dimension (*N* = 49)

To simulate emergent tissue behaviour, the lattice must satisfy two criteria: (1) statistically sufficient representation of the heterogeneity-derived pacemaker population, and (2) minimisation of boundary artefacts. We define the **Core** as cells with 4 neighbours and the **Boundary** as cells with *<* 4 neighbours.

- **Case** *N* = 25 (5 × 5): Core = 9 cells, Boundary = 16 cells. The boundary comprises 64% of the tissue. Since boundary cells lack current sinks on at least one side, they are artificially excitable. In this configuration, spontaneous activity is statistically more likely to arise from a boundary artefact than from a true internal pacemaker.
- **Case** *N* = 49 (7×7): Core = 25 cells, Boundary = 24 cells. This is the smallest regular square lattice where the Core (*>* 50%) statistically dominates.

Given the pacemaker probability density function (*P*_*pacer*_ ≈ 3 − 15%), a 7 × 7 grid yields an expected pacemaker population of ≈ 4 − 8 cells. This size ensures that the pacemaker clusters are statistically likely to reside within the core, permitting the observation of true radial wave propagation uncorrupted by edge effects.

